# Glutamate co-release from catecholaminergic neurons shapes breathing and is inhibited during opioid-induced respiratory depression

**DOI:** 10.64898/2026.03.25.714271

**Authors:** Alexis Riley-DiPaolo, Valerie V. Cabrera, Umit M. Akkaya, Sebastian N. Maletz, Adrienn G. Varga

## Abstract

Breathing is controlled by a distributed brainstem network that includes multiple catecholaminergic nuclei. The locus coeruleus (LC), the brain’s primary source of noradrenaline (NA), projects to several respiratory centers, including the Kölliker-Fuse (KF) nucleus in the pons. While LC neurons are predominantly noradrenergic (NAergic), many co-release glutamate, which may contribute to the state-dependent modulation of breathing, particularly during opioid exposure. Here, we examined how opioids affect NAergic and glutamatergic signaling in the LC-KF circuit using optogenetics and whole-cell patch clamp recordings in mouse brain slices. Optogenetic activation of LC terminals evoked glutamatergic excitatory postsynaptic currents (EPSCs) in KF neurons that were presynaptically inhibited by the opioid receptor agonist Met-enkephalin. Additionally, ∼36% of glutamate-responsive KF neurons exhibited postsynaptic opioid inhibition via GIRK currents, while KF neurons receiving excitatory NAergic input showed minimal opioid sensitivity. To assess the behavioral role of glutamate release from all catecholaminergic neurons, we compared breathing in awake VGluT2fl/fl::TH-Cre mice (lacking VGluT2 in tyrosine hydroxylase-positive neurons) to control littermates and TH-Cre hemizygous mice using whole-body plethysmography. The conditional VGluT2 knockout mice exhibited prolonged inspiratory duration, increased tidal volume, and reduced respiratory rate during baseline breathing, with state-dependent differences emerging during hypercapnia. Systemic morphine administration diminished these genotype differences, and machine learning analysis using dynamic time warping confirmed that genotype-specific breathing patterns were distinguishable at baseline, but not after morphine. These findings demonstrate that glutamate co-release from catecholaminergic neurons modulates respiratory patterning in a state-dependent manner and is selectively vulnerable to opioid inhibition.

## Introduction

Breathing is orchestrated by a distributed network of medullary and pontine brainstem nuclei (1, 2). Within this network, catecholaminergic neurons provide homeostatic and state-dependent control of respiratory output (2–5). The physiological importance of catecholaminergic modulation is evidenced by multiple disorders with respiratory symptoms linked to catecholaminergic dysfunction, including Rett syndrome, sudden infant death syndrome, and congenital central hypoventilation syndrome (6–10). Targeted chemogenetic inhibition of catecholaminergic neurons disrupts respiratory chemoreflexes and metabolic homeostasis (11), confirming their necessity for appropriate respiratory responses to metabolic challenges. However, the contributions of secondary signaling mechanisms (non-catecholamine release) and the role of specific catecholaminergic nuclei remain incompletely understood.

The locus coeruleus (LC) is the brain’s most broadly connected catecholaminergic nucleus and the primary source of central noradrenaline (NA) (12–16). Beyond its extensive mid- and forebrain projections, the LC innervates key respiratory nuclei including the pre-Bötzinger complex, Bötzinger complex, nucleus tractus solitarius, parabrachial complex, and Kölliker-Fuse (KF) nucleus (2, 15, 17–19). The LC influences respiratory homeostasis through multiple mechanisms. The LC contributes to central chemosensation and state-dependent control of breathing (12, 20). More specifically, activation of *Phox2b* expressing LC neurons has been shown to increase basal ventilation in awake mice (21–23).

While the majority of LC neurons are primarily noradrenergic (NAergic), many co-release glutamate, dopamine and various neuropeptides (24–28). Recent evidence demonstrates that LC neurons send monosynaptic glutamatergic projections to the KF (18), supporting neurotransmitter multiplexing where glutamate and NA provide distinct, but coordinated, regulatory signals. Interestingly, glutamate release from central NAergic neurons (identified using dopamine beta-hydroxylase [DBH]-Cre mice) does not critically influence baseline breathing or chemosensory reflexes (29), suggesting a more specific, state-dependent role for this channel of excitatory signaling.

One such state-dependent challenge is opioid-induced respiratory depression (OIRD), which results from mu-opioid receptor (MOR) activation throughout the central nervous system (30, 31). The KF is critical for opioid-induced respiratory rate depression and pattern irregularity (32–34). Notably, MORs are highly abundant in both LC neurons (>80%) (35) and KF neurons (∼60%) (32), positioning the LC-KF circuit as a potentially important pathway for OIRD. However, how catecholaminergic systems, individually and in general, support breathing under opioid exposure remains unclear.

Here, we relied on the LC-KF circuit as an *ex vivo* model to investigate the opioid sensitivity of NAergic and glutamatergic signaling using optogenetics and whole-cell patch clamp recordings. We also examined whether glutamate release from all catecholaminergic (tyrosine hydroxylase [TH]-positive) neurons throughout the brain contributes to baseline breathing and OIRD using conditional VGluT2 knockout mice and whole-body plethysmography. Our results reveal that LC glutamatergic signaling is vulnerable to both presynaptic and postsynaptic opioid inhibition, while NAergic excitation appears resistant to both. Furthermore, removal of VGluT2 from catecholaminergic neurons alters respiratory patterning in a state-dependent manner. These glutamatergic influences are diminished by morphine administration, consistent with our LC-KF circuit-specific findings of preferential opioid sensitivity in the glutamatergic component of this system.

## Methods

### Animals

Adult mice (male and female, 3-6 months old) were used for all experiments. Mice were bred and maintained in-house at the University of Florida. Mice were group-housed in standard sized plastic cages and kept on a 12 h light–dark cycle, with water and food available *ad libitum*. All procedures conducted were in accordance with the National Institutes of Health guidelines and with approval from the Institutional Animal Care and Use Committee of the University of Florida. TH-Cre (B6.Cg-7630403G23Rik^Tg^ (Th–cre)^1Tmd^/J JAX#: 008601) hemizygous mice (referred to as TH-Cre in this manuscript) were used for brain slice recording and optogenetic experiments. To eliminate the ability of TH+ neurons to release glutamate, TH-Cre mice were crossed with VGluT2^fl/fl^ mice (Slc17a6^tm1Lowl^/J JAX#: 012898) to create VGluT2^fl/fl^::TH-Cre mice that were used for whole-body plethysmography.

### Stereotaxic injections

Mice were anesthetized with isoflurane (2-4% in 100% oxygen, Zoetis) and mounted in a stereotaxic frame (Kopf Instruments, Tujunga, CA) where anesthesia was maintained through a nose cone. Body temperature was maintained with a water circulating heating pad (37⁰C). Meloxicam (5mg/kg, s.c., Patterson Veterinary) and lidocaine (2mg/kg s.c., Patterson Veterinary) were administered before midline incision and exposure of the skull. The dorsal skull was levelled horizontally between bregma and lambda. A small craniotomy was made to access the LC (y = -5.4 mm, x = ± 0.85 mm) bilaterally. A glass micropipette was filled with AAV5-EF1α-DIO-hChR2(H134R)-EYFP-WPRE-HGHpA (Addgene, 202298-AAV5, titer 1×10¹³ vg/mL), then lowered into the LC (z = -3.65 mm) and the virus (250 nl/ hemisphere) was injected bilaterally with a Nanoject III injector (Drummond Scientific Company, Broomall, PA, USA) at 50nl/sec in 10nl increments with 20 sec inter-pulse intervals. Following each injection, the pipette was left in place for ∼10 min and slowly retracted over the course of at least 5 min. The wound was closed with Vetbond tissue adhesive (3M Animal Care Products, St. Paul, MN, USA). Mice were then placed in a recovery chamber and kept warm until they ambulated normally. Slice electrophysiology experiments were performed 6-10 weeks later. Placement of injections was subsequently verified in brain slices (50-100 µm fixed or 230 µm live slice) from every mouse by visualizing eYFP fluorescence using a multi-zoom microscope (Nikon AZ100).

### Brain slice electrophysiology

Mice were deeply anesthetized with isoflurane and decapitated. The brain was quickly removed and mounted in a vibratome chamber (Leica VT 1200S, Leica Biosystems, Buffalo Grove, IL, USA). Coronal slices (230 µm) containing the LC or KF (identified based on anatomical markers and coordinates from Paxinos and Franklin (36) were cut in warm (34°C), carbogenated artificial cerebrospinal fluid (ACSF) that contained the following (in mM): 126 NaCl, 2.5 KCl, 1.2 MgCl_2_, 2.4 CaCl_2_, 1.2 NaH_2_PO_4_, 11 D-glucose and 21.4 NaHCO_3_ (equilibrated with 95% O_2_/5% CO_2_) (36). Slices were stored at 32°C in glass vials with equilibrated ACSF. MK801 (10 µM) was added to the cutting solution and for the initial incubation of slices (30-60 min) to prevent NMDA receptor-mediated excitotoxicity. After incubation, the slices were transferred to a recording chamber that was perfused with 34°C ACSF at a rate of 1.5-3 ml/min. The temperature of the solution was maintained with an in-line closed-loop heater (Warner Instruments/ Harvard Apparatus, Holliston, MA).

Cells were visualized using an upright microscope (Nikon FN1) equipped with custom built IR-Dodt gradient contrast illumination. For recordings performed in the LC, eYFP fluorescence was identified in infected LC cells using LED epifluorescent illumination, eYFP excitation/emission filter cube and detected using a DAGE-MTI IR-1000 camera with sufficient sensitivity in the eYFP emission range. To avoid neurotransmitter depletion in the LC axon terminals located in KF slices, recordings from postsynaptic KF neurons were done without fluorescence visualization.

Whole-cell recordings were performed with a Multiclamp 700B amplifier (Molecular Devices, Sunnyvale, CA) in voltage-clamp (V_hold_= -60mV) or current clamp mode. Recording pipettes (1.5 – 3 MΩ) were filled with internal solution containing (in mM): 115 potassium methanesulfonate, 20 NaCl, 1.5 MgCl_2_, 5 HEPES(K), 2 BAPTA, 1-2 Mg-ATP, 0.2 Na-GTP, adjusted to pH 7.35 and 275-285 mOsM. Liquid junction potential (10 mV) was not corrected. Data were filtered at 10 kHz and collected at 20 kHz with pClamp10.7 (Molecular Devices, Sunnyvale, CA) and simultaneously collected at 400 Hz with PowerLab (Lab Chart version 8; AD Instruments, Colorado Springs, CO). Series resistance was monitored without compensation and remained < 20 MΩ for inclusion in data processing and analysis. To block inhibitory fast synaptic transmission in KF recordings, picrotoxin (100µM) was added to the ACSF. Additional drugs listed in specific experiments were applied by bath perfusion at the indicated concentrations.

Wide-field stimulation of Channel-rhodopsin2 (ChR2) with blue light was achieved with a 470nm LED (1.0A, Thorlabs) controlled by a Thorlabs T-Cube TEC controller and directed through a YFP filter cube and the 40x objective, which produced 8.5 mW/cm^2^ at the slice, confirmed by measurements with a photodiode. To verify efficient functional expression of ChR2 in LC neurons, whole-cell voltage clamp recordings were performed from eYFP-labeled cells in the LC in brainstem slices and stimulated with blue light at increasing intensities while the cell was held at -60 mV (10%, 25%, 100% of max intensity, 100ms light pulse) (18). In current clamp, the ability to evoke action potentials with a 100pA current injection (200ms) was compared to a 200ms blue light pulse to verify that optical stimulation induces firing in ChR2-eYFP expressing LC neurons. Control experiments were performed on non-eYFP labeled LC neurons in the same slice, which did not display evoked currents or action potentials.

LC axon terminals expressing ChR2 were optically stimulated in KF slices during whole-cell recordings of KF neurons. Wide field blue light stimulation (470 nm, 10 sec continuous) was used to achieve presynaptic NA release and to measure persistent NAergic currents in postsynaptic KF neurons. Light-evoked persistent current amplitude was measured in Lab Chart 8 (AD Instruments, Colorado Springs, CO). First, traces were smoothed with a Bartlett window (7-sample sliding window), and the peak amplitude was defined as the maximum value. The maximum value was then subtracted from the mean amplitude of a 30-60 sec baseline immediately preceding the onset of the current. For inclusion in further testing and final analysis, persistent currents had to be reproduced at least twice.

The glutamate release-stimulating protocol involved 2×5 ms paired light pulses with a 50 ms inter-stimulus interval, repeated every 20 seconds at least 5 times. Glutamatergic events were analyzed in Clampfit 10.7 (Molecular Devices, Sunnyvale, CA). Each light-evoked oEPSC episode in a sweep (typically 10–50 sweeps per condition) was digitally filtered at 1 kHz with a Gaussian window, then adjusted to baseline immediately before light onset. Only recordings with qualities indicating monosynaptic glutamate-release (latency < 6ms, low SD of latencies or “jitter” < 2ms) were used for analysis. Single sweep values were averaged together to obtain a mean oEPSC amplitude for each condition. Average oEPSC amplitudes across conditions were then compared to each other.

### Drugs

All drugs were reconstituted and stored according to the manufacturers’ directions. (+)-MK801 hydrogen maleate, picrotoxin, ME ([Met5]-enkephalin acetate salt), bestatin HCl, DL-thiorphan were obtained from Sigma-Aldrich (St. Louis, MO).

Morphine sulfate was obtained from the National Institute on Drug Abuse Supply Program (RTI International, Research Triangle Park, NC).

### Whole-body plethysmography

Whole-body plethysmography was used to measure the respiration of awake, freely moving mice as described previously (37). Recordings of respiratory frequency and estimated tidal volume (weight adjusted post-hoc) were collected using vivoFlow whole-body plethysmography and IOX2 software (SCIREQ Inc, Montreal, QC, Canada). Across all groups, mice were acclimated to the plethysmography chamber ventilated with standard air (0.5 L/min) for 1 hour / day on three consecutive days prior to testing to minimize possible influences of stress or exploration on respiratory measurements. On testing day, mice were placed into the plethysmograph chambers, where respiration was measured using a pressure transducer. Recording conditions included a 30-minute period of normoxia with standard air (21% O_2_, 79% N_2_), followed by a 15-minute hypercapnic challenge (21% O_2_, 7% CO_2_, 72% N_2_), then a 10-minute period of standard air prior to morphine injection. Following morphine administration (30 mg/kg in saline, i.p.), mice underwent a 20-minute period of normoxia (referred to as normoxia + morphine) and another 15-minute hypercapnic challenge (referred to as hypercapnia + morphine) to assess morphine-induced respiratory depression and any opioid-induced blunting of the hypercapnic ventilatory response before completion of the experiment.

### Statistical Analysis

All statistical analyses were performed in Prism v10 (GraphPad, Boston, MA). All error bars represent SEM unless otherwise stated. Data with n > 8 were tested for normality with Kolmogorov-Smirnov test. Normally distributed data were analyzed with parametric tests, and non-normally distributed data were analyzed with non-parametric tests. Specific statistical tests and *p* values can be found in the tables or results.

### Machine learning to classify breathing

Whole-body plethysmography data were further analyzed with the k-Nearest Neighbors with Dynamic Time Warping (DTW-kNN) machine learning algorithm (38). This classic, shape-based classifier was used for measuring the similarity between the time series respiratory frequency data obtained from the three groups of mice. For each subject, two separate time-series recordings of instantaneous respiratory frequency were obtained (sampled at 100 Hz) and underwent Z-normalization to ensure uniform scale across all features. The time series were then further segmented into 1 second samples resulting in a total of 7200 samples for normoxia baseline and 7200 samples for the normoxia + morphine condition. Subsequently, we employed a 3-Fold Group Cross-Validation scheme for training and testing (GroupKFold in scikit-learn). The model accuracy reported is the average performance across 3 folds. The performance of the model is also evaluated based on its F1-score.

DTW was used to find the optimal non-linear alignment between two samples by warping the time axes, allowing for comparisons between sequences with different temporal structures (39). The classifier then creates a cost matrix, where each value represents the dissimilarity (or distance) between corresponding points in the compared time series. The kNN algorithm was used for classification of unknown samples based on the majority class of its k-nearest neighbors. The model was configured with k=3, where classification is determined by the majority vote of 3 neighbors, that represent the lowest DTW distance (i.e. highest morphological similarity) in the training set.

All data pre-processing, analysis, and model validation were conducted within a Google Colaboratory environment (Google Brain Team, Mountain View, CA, USA). The analysis was implemented in Python (v3.12), utilizing a suite of open-source scientific computing libraries. Key libraries included: pandas for data manipulation, numpy for numerical operations, scikit-learn for core machine learning utilities (including GroupKFold and LabelEncoder), and tslearn (v0.6.4) for the dtw distance metric (40).

## Results

### The LC-KF circuit is both presynaptically and postsynaptically inhibited by opioids

To begin to elucidate the contribution of NA and glutamate release in the context of OIRD, we first tested the pre- and postsynaptic impact of opioids in the specific LC-KF circuit. We selectively expressed ChR2 bilaterally in the LC NAergic population of TH-Cre mice and performed whole-cell voltage clamp recordings in brain slices that contain KF neurons and LC axon terminals (**Fig 1A**) (18). Short blue light pulses (5ms) elicited optically-evoked glutamatergic excitatory postsynaptic currents (oEPSCs) that were AMPA-receptor mediated (**Fig 1B**, shown in more detail in Varga et al., 2025). We recorded from each KF neuron while delivering paired light pulses (5ms each, 50ms inter-pulse interval, 20 sec inter-sweep-interval) in baseline conditions and during bath application of the opioid receptor agonist Met-enkephalin (ME, 1 µM). Across all recordings (n = 11 neurons from n = 9 mice) bath application of ME led to a significant decrease in the amplitude of the first oEPSC (mean baseline amplitude ± SEM: 77.49pA ± 20.74pA vs. mean ME amplitude ± SEM: 59.97pA ± 16.19pA, *p* = 0.042 by Wilcoxon matched-pairs signed rank test; **Fig 1C**), indicating presynaptic inhibition of glutamate release from LC terminals onto KF neurons, as expected based on the high abundance of MORs on LC neurons (41–43). The temporal dynamics of glutamate release remained similar however, resulting in no statistical difference in latency (mean baseline latency ± SEM: 3.13ms ± 0.43ms vs. mean ME latency ± SEM: 3.19ms ± 0.39ms, *p* = 0.571 by two-tailed paired *t* test; **Fig 1D**) and no change in the consistency of oEPSC latency or jitter following opioid administration in the bath (mean SD in oEPSC latency during baseline ± SEM: 0.782ms ± 0.136ms vs. during ME ± SEM: 0.727ms ± 0.135ms; *p* = 0.570 by two-tailed paired *t* test; **Fig 1E**). Next, we assessed the influence of ME on the probability of vesicle release by analyzing the paired pulse ratio (PPR) of the second postsynaptic current amplitude over the first (**Fig 1G**). Out of the 11 neurons, 7 showed a significant increase in PPR, providing additional evidence for opioid-induced presynaptic inhibition in this circuit (mean PPR during baseline ± SEM: 0.779 ± 0.128 vs. mean PPR during ME ± SEM: 0.935 ± 0.139; *p* = 0.044 by two-tailed paired *t* test).

**Figure 1:**
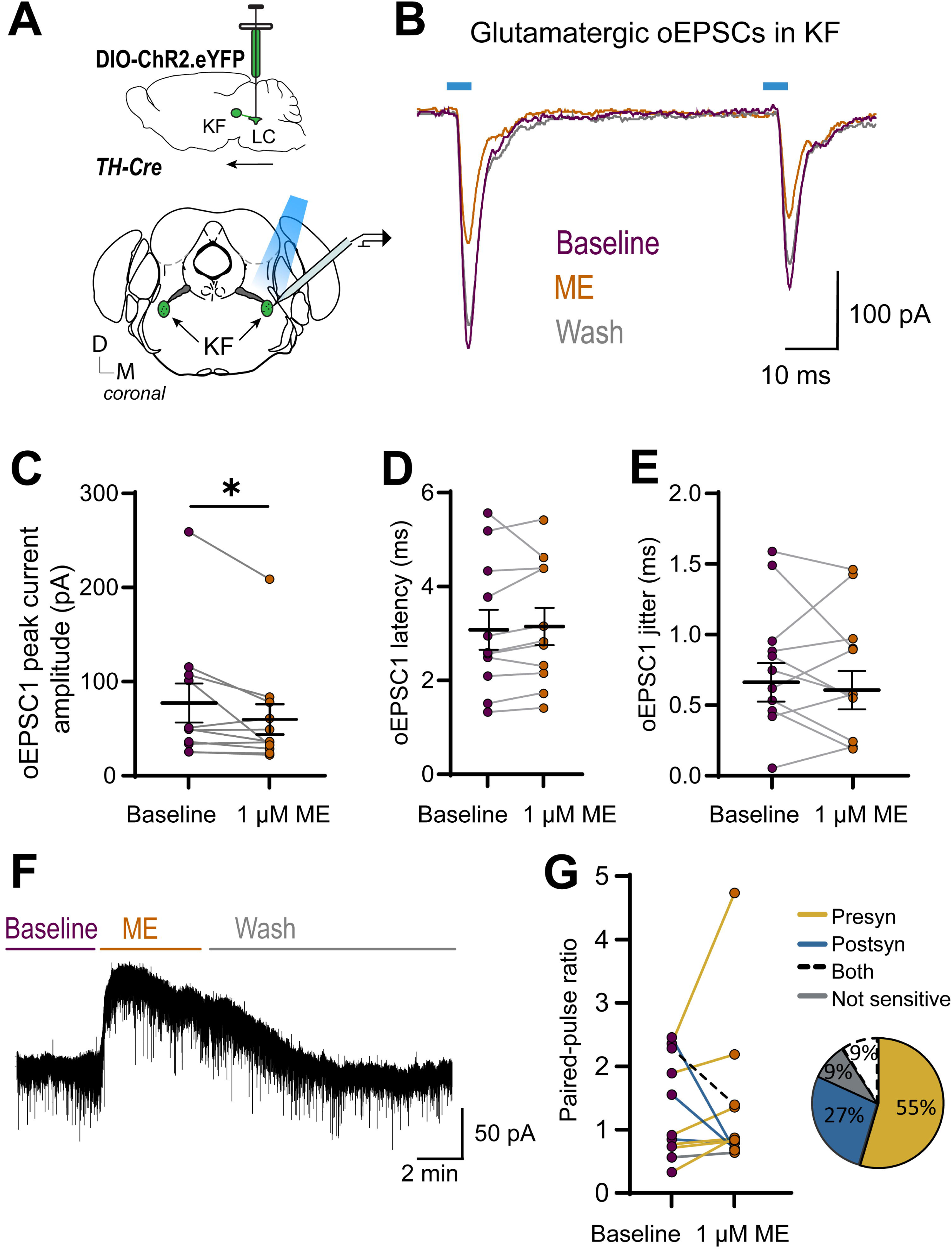
Synaptic opioid inhibition in the LC and KF circuit. TH-Cre mice with selectively expressed ChR2 bilaterally in the LC were used for whole-cell voltage clamp recordings in slices containing KF neurons and LC axon terminals. (**A**) Experimental schematic. (**B**) Light pulses elicited AMPA-receptor mediated oEPSCs. ME exposure led to (**C**) a significant decrease in amplitude of first oEPSC, (**D**) no difference in latency, (**E**) or jitter. (**F**) Representative GIRK current in the KF. (**G**) 55% of neurons displayed a significant increase in PPR following ME administration.

The direct, somatodendritic effects of opioids on ∼60% of KF neurons are well-established (32), however, it is not known what proportion of LC glutamate- or NA-modulated KF neurons are opioid-sensitive. Here, we found that bath application of ME results in a reversible and reproducible G protein-coupled inwardly rectifying potassium channel current (GIRK) (31, 32) in 4 out of the 11 neurons indicating postsynaptic opioid-inhibition in a portion of LC glutamate-modulated KF cells (**Fig 1F** example GIRK current in KF, **Fig 1G**).

Given that, in addition to glutamate, α1-receptor activation by NA binding may also excite 45% of KF neurons, we investigated the opioid-sensitivity of KF neurons that display persistent excitatory NAergic currents in response to 10 sec blue light stimulation (18). We did not find any presynaptic inhibitory effects of the opioid agonist ME (3 µM) on NA release in 6 out of 6 cells (n=4 mice; mean baseline NA current amplitude ± SEM: 11.43pA ± 1.88pA vs. during 3 µM ME ± SEM: 10.06pA ± 2.67pA; *p* = 0.458 by two-tailed paired *t* test). However, this is likely due to the nature of the slice preparation, where KF containing slices only encompass LC axon terminals, and no cell bodies where MORs may be located.

Additionally, out of the 8 KF neurons (from n= 6 mice) that had an excitatory α1-receptor mediated persistent current, only 1 neuron had a GIRK current in response to ME, suggesting that the majority of KF neurons excited by NA are not directly opioid-sensitive.

### VGluT2-based glutamatergic signaling in catecholaminergic neurons contributes to the control of breathing in a state-dependent manner

Catecholaminergic (TH-positive) neurons are necessary for the control of some aspects of baseline breathing, as well as chemosensory responses elicited by hypercapnic and hypoxic conditions (11, 44, 45). However, glutamate release from central NAergic neurons (those containing dopamine beta hydroxylase [DBH], in this case in DBH-Cre mice) does not appear to have an impact on baseline, hypercapnic, or hypoxic breathing (29). It remains unknown whether glutamatergic signaling from the broader population of catecholaminergic (TH-expressing) neurons contributes to respiratory control, and how this potential influence may be attenuated by opioids is also unclear.

To fill this gap, we compared the breathing of freely moving VGluT2^fl/fl^::TH-Cre mice (n=4) that lack the machinery that supports glutamate release from TH+ neurons, to TH-Cre (n=4) and littermate VGluT2^f/fl^::WT mice (n=4) using whole-body plethysmography (**Fig 2 and Tables 1-2**). As described above, all three groups were first tested under normoxic conditions to assess baseline breathing (see example traces in **Fig 2A, B, C**), then exposed to 20 minutes of normoxic hypercapnia (21% O_2_/7% CO_2_), followed by normoxia (21% O_2_) to allow breathing to return to baseline. Following this, mice received a systemic injection of a moderately respiratory depressant dose of morphine (30 mg/kg i.p., in saline), then returned to the plethysmograph chamber for 20 minutes of normoxia and 15 minutes of hypercapnia to assess the influence of morphine on breathing.

**Figure 2:**
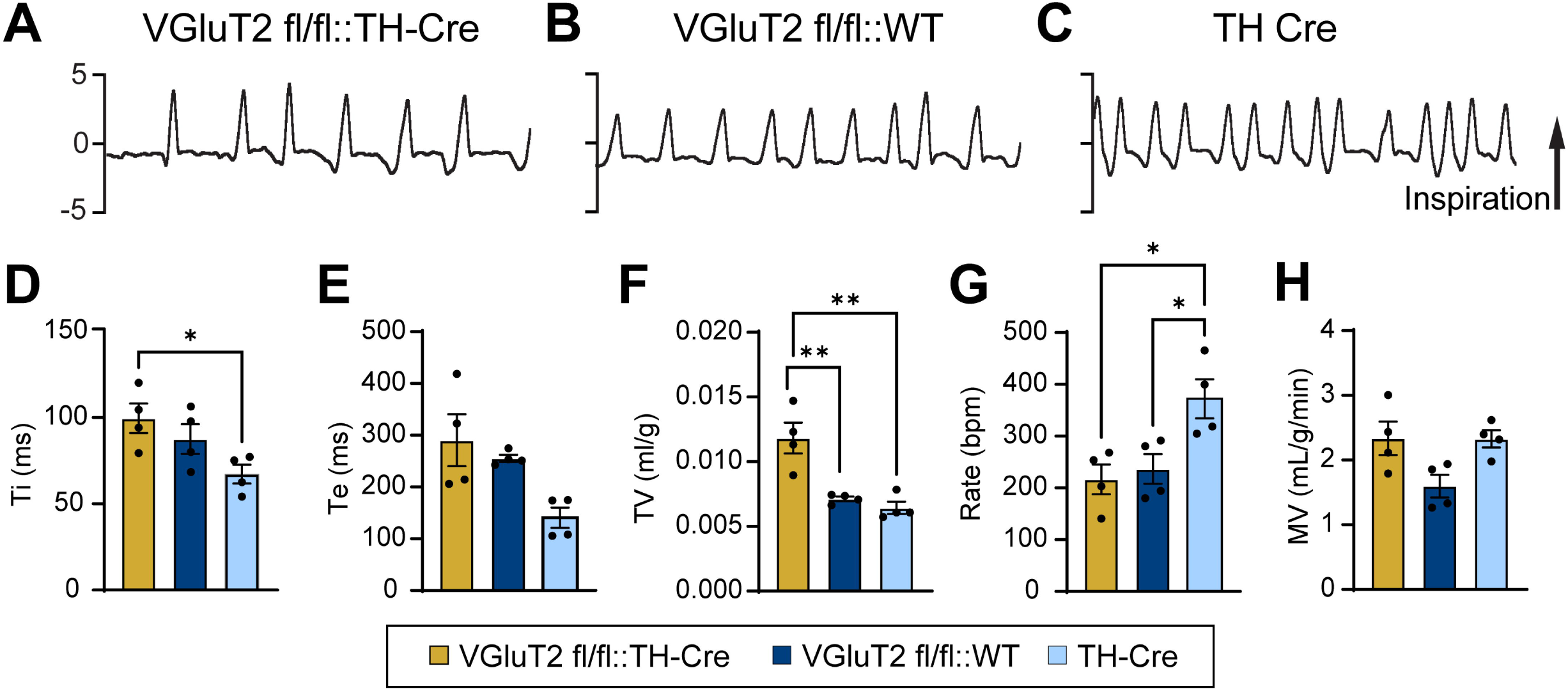
VGluT2 deficient mice exhibit alterations in basal breathing. (**A**)-(**C**) are representative whole-body plethysmograph traces (4 sec duration). Animals were tested in whole-body plethysmography under normoxic conditions. Analysis revealed (**D**) a significant increase in inspiratory time (Ti), (**E**) no change in expiratory time (Te), (**F**) an increase in tidal volume (TV), (**G**) a decrease in respiratory rate, and (**H**) no change in minute ventilation (MV) from VGluT2 deficient animals.

**Table 1:**
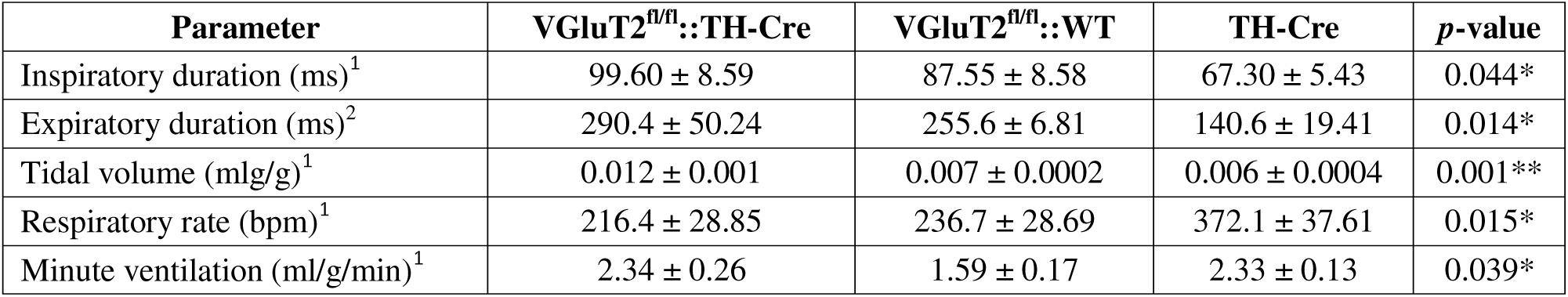
Summary of respiratory parameters by all groups during normoxia (Fig 2) . ^1^One-way ANOVA (parametric); ^2^Kruskal-Wallis test (nonparametric); \**p* < 0.05; \*\**p* < 0.01. Values are reported as mean ± SEM unless otherwise stated.

**Table 2:** Summary of between-group comparisons during normoxia (Fig 2). ^3^Tukey’s multiple comparisons test (parametric); ^4^ Dunn’s multiple comparisons test (nonparametric); \**p* < .05; \*\**p* < .01.

| Parameter $p$-value | VGluT2<sup>fl/fl</sup>::TH-Cre vs.<br>VGluT2<sup>fl/fl</sup>::WT | VGluT2<sup>fl/fl</sup>::TH-Cre vs.<br>TH-Cre | VGluT2<sup>fl/fl</sup>::WT vs.<br>TH-Cre |
| --- | --- | --- | --- |
| Inspiratory time<sup>3</sup> | 0.53 | **0.037\*** | 0.2 |
| Expiratory time<sup>4</sup> | 0.99 | 0.055 | 0.055 |
| Tidal volume<sup>3</sup> | **0.0041\*\*** | **0.0017\*\*** | 0.79 |
| Respiratory rate<sup>3</sup> | 0.89 | **0.018\*** | **0.036\*** |
| Minute ventilation<sup>3</sup> | 0.06 | 0.99 | 0.06 |

We found that in normoxic, baseline conditions mice without VGluT2 in catecholaminergic cells have significantly longer inspiratory durations (**Fig 2D**), no statistical difference in expiratory durations (**Fig 2E**), significantly larger tidal volumes (**Fig 2F**), and significantly lower respiratory rates compared to TH-Cre controls (**Fig 2G**). When compared to the VGluT2^fl/fl^::WT group, only tidal volume was statistically different (**Fig 2**). Overall, there was a statistically significant effect of genotype on minute ventilation, but individual comparisons between the three groups were not significant (**Fig 2H**), indicating that the reciprocal difference in tidal volume and respiratory rate produces similar overall minute ventilation, which consists of slower, larger breaths for the mice lacking glutamate in TH+ neurons, and faster, smaller breaths in the control TH-Cre animals.

Next, we examined how glutamate signaling contributes to the hypercapnic ventilatory response, and whether it is diminished during morphine-induced respiratory depression and simultaneous hypercapnic challenge (**Fig 3**, **Tables 3-4**). Subjects in all groups responded to hypercapnia with an increase in rate, tidal volume, and minute ventilation, a decrease in expiratory duration and no substantial change in inspiratory duration, as expected (**Fig 3**, 7% CO_2_ columns). Similarly, morphine decreased respiratory rate and tidal volume in all groups, which corresponded with increases in both inspiratory and expiratory durations, and an overall decrease in minute ventilation (**Fig 3**, morphine columns). Exposing the mice to a hypercapnic challenge during morphine-induced respiratory depression brought all breathing parameters closer to baseline levels, indicating that the hypercapnic ventilatory reflex is only minimally blunted at this level of OIRD (**Fig 3**, 7% CO_2_ with morphine columns).

**Figure 3:**
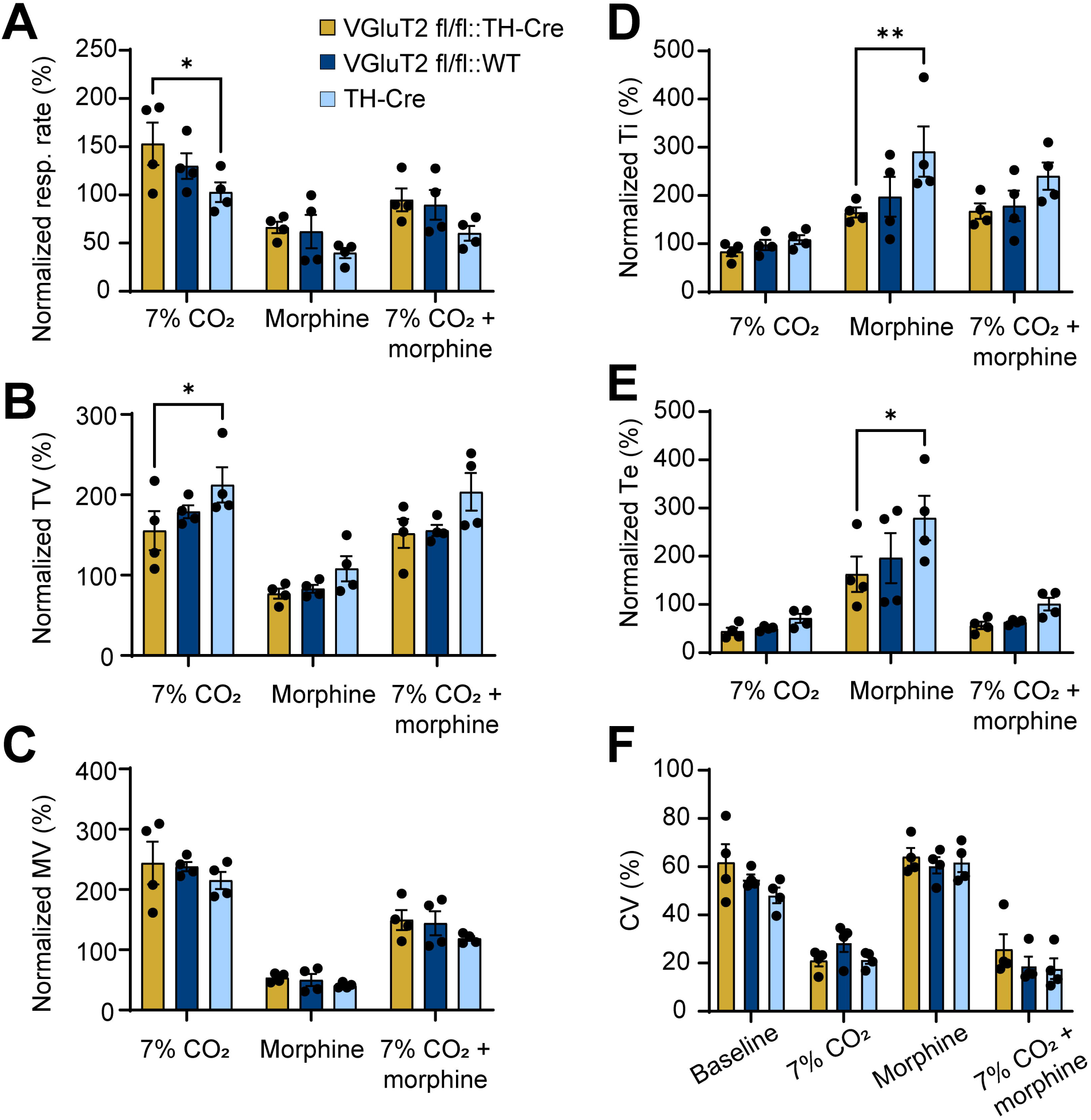
VGluT2 conditional knockout drives nuanced changes in breathing during hypercapnia and following morphine exposure. Animals were tested in whole-body plethysmography under normoxic and hypercapnic conditions, before and after morphine administration. Respiratory outputs are normalized to baseline breathing during normoxia. Analysis revealed (**A**) an increase in hypercapnic breathing rate, (**B**) a decrease in tidal volume (TV), (**C**) no change in minute ventilation (MV), (**D**) an increase in inspiratory time (Ti) following morphine exposure, (**E**) a decrease in expiratory time (Te), (**F**) and no change in coefficient of variation (CV) of respiratory rate from VGluT2 deficient animals

**Table 3:** Summary of respiratory parameters between groups (hypercapnia/morphine) (Fig 3). Data were normalized to baseline normoxic breathing.

| Parameter (% of baseline) | Condition | VGluT2<sup>fl/fl</sup>::TH-Cre | VGluT2<sup>fl/fl</sup>::WT | TH-Cre |
| --- | --- | --- | --- | --- |
| Ti | 7% CO<sub>2</sub> | 83.452 ± 9.134 | 97.585 ± 10.563 | 108.455 ± 9.227 |
|  | Normoxia + morphine | 164.905 ± 10.727 | 197.213 ± 41.246 | 291.052 ± 52.167 |
|  | 7% CO<sub>2</sub> + morphine | 167.727 ± 16.066 | 178.658 ± 31.645 | 240.195 ± 28.273 |
| Te | 7% CO<sub>2</sub> | 44.221 ± 7.770 | 50.600 ± 2.368 | 71.440 ± 9.384 |
|  | Normoxia + morphine | 162.551 ± 36.675 | 196.330 ± 51.862 | 279.360 ± 46.079 |
|  | 7% CO<sub>2</sub> + morphine | 56.625 ± 7.567 | 63.419 ± 2.494 | 100.934 ± 13.221 |
| TV | 7% CO<sub>2</sub> | 155.314 ± 24.212 | 178.814 ± 7.989 | 212.436 ± 21.855 |
|  | Normoxia + morphine | 77.060 ± 6.228 | 83.052 ± 4.878 | 108.031 ± 15.656 |
|  | 7% CO<sub>2</sub> + morphine | 151.916 ± 17.909 | 155.659 ± 6.885 | 203.648 ± 23.393 |
| Rate | 7% CO<sub>2</sub> | 152.941 ± 21.919 | 129.960 ± 13.333 | 102.854 ± 10.237 |
|  | Normoxia + morphine | 66.298 ± 5.973 | 61.881 ± 17.286 | 39.813 ± 5.296 |
|  | 7% CO<sub>2</sub> + morphine | 94.896 ± 11.880 | 89.710 ± 15.457 | 60.269 ± 7.586 |
| MV | 7% CO<sub>2</sub> | 243.916 ± 35.564 | 237.856 ± 7.765 | 215.025 ± 14.079 |
|  | Normoxia + morphine | 53.452 ± 3.711 | 49.893 ± 9.929 | 40.869 ± 2.278 |
|  | 7% CO<sub>2</sub> + morphine | 149.462 ± 16.492 | 144.058 ± 19.886 | 118.677 ± 3.973 |

**Table 4:** Summary of respiratory parameters between groups (hypercapnia/morphine) (Fig 3). Data were normalized to baseline normoxic breathing. Tukey’s multiple comparisons test (parametric); \**p* < 0.05; \*\**p* < 0.01.

| Parameter,<br>*p*-value | Condition | VGluT2<sup>fl/fl</sup>::TH-Cre<br>vs.<br>TH-Cre | VGluT2<sup>fl/fl</sup>::TH-Cre<br>vs.<br>VGluT2<sup>fl/fl</sup>::WT | VGluT2<sup>fl/fl</sup>::WT<br>vs.<br>TH-Cre |
| --- | --- | --- | --- | --- |
| Ti | 7% CO<sub>2</sub> | 0.7997 | 0.9307 | 0.9583 |
|  | Normoxia + morphine | **0.0089\*\*** | 0.6901 | 0.0592 |
|  | 7% CO<sub>2</sub> + morphine | 0.1717 | 0.9579 | 0.2738 |
| Te | 7% CO<sub>2</sub> | 0.7577 | 0.9847 | 0.8493 |
|  | Normoxia + morphine | **0.0133\*** | 0.654 | 0.0936 |
|  | 7% CO<sub>2</sub> + morphine | 0.4858 | 0.9827 | 0.5935 |
| TV | 7% CO<sub>2</sub> | **0.0479\*** | 0.5659 | 0.3203 |
|  | Normoxia + morphine | 0.3781 | 0.9629 | 0.5266 |
|  | 7% CO<sub>2</sub> + morphine | 0.0786 | 0.9853 | 0.1088 |
| Rate | 7% CO<sub>2</sub> | **0.0316\*** | 0.4445 | 0.3283 |
|  | Normoxia + morphine | 0.3445 | 0.9695 | 0.4726 |
|  | 7% CO<sub>2</sub> + morphine | 0.1704 | 0.9583 | 0.2715 |
| MV | 7% CO<sub>2</sub> | 0.4231 | 0.9616 | 0.5803 |
|  | Normoxia + morphine | 0.8455 | 0.9866 | 0.917 |
|  | 7% CO<sub>2</sub> + morphine | 0.3783 | 0.9694 | 0.5122 |

The results also revealed genotype-dependent differences that support the recruitment of glutamate release in the state-dependent control of breathing. During hypercapnia, VGluT2^fl/fl^::TH-Cre mice had increased respiratory rates (**Fig 3A**), and smaller tidal volumes (**Fig 3B**) as compared to TH-Cre mice. Following morphine administration, VGluT2^fl/fl^::TH-Cre mice had shorter inspiratory (**Fig 3D**) and expiratory durations (**Fig 3E**), compared to TH-Cre mice. However, there was ultimately no difference in overall minute ventilation across all conditions (no main effect of genotype two-way rmANOVA F(2,27) = 1.88, *p* = 0.173; **Fig 3C**). Finally, we did not find any evidence of glutamatergic influence on respiratory pattern variability as measured by the coefficient of variation (CV%) analysis of instantaneous breathing frequency in baseline, hypercapnia, or morphine, and hypercapnia with morphine trials (no main effect of genotype by two-way rmANOVA F(2,36) = 1.99, *p* = 0.15; **Fig 3F**).

### Machine learning reveals VGluT2-dependent breathing patterns during basal breathing, that are eliminated by opioid-induced respiratory depression

While traditional parameter analyses revealed some specific differences between genotypes under different metabolic states, we applied the k-Nearest Neighbors with Dynamic Time Warping (DTW-kNN) classifier to determine whether the overall temporal structure of breathing patterns could distinguish genotypes (visualized comparison of two example samples from TH-Cre mice with DTW, **Fig 4A**). This approach integrates information across multiple breath cycles and captures intricate dynamics in breathing rhythm that single-parameter comparisons may not detect.

**Figure 4:**
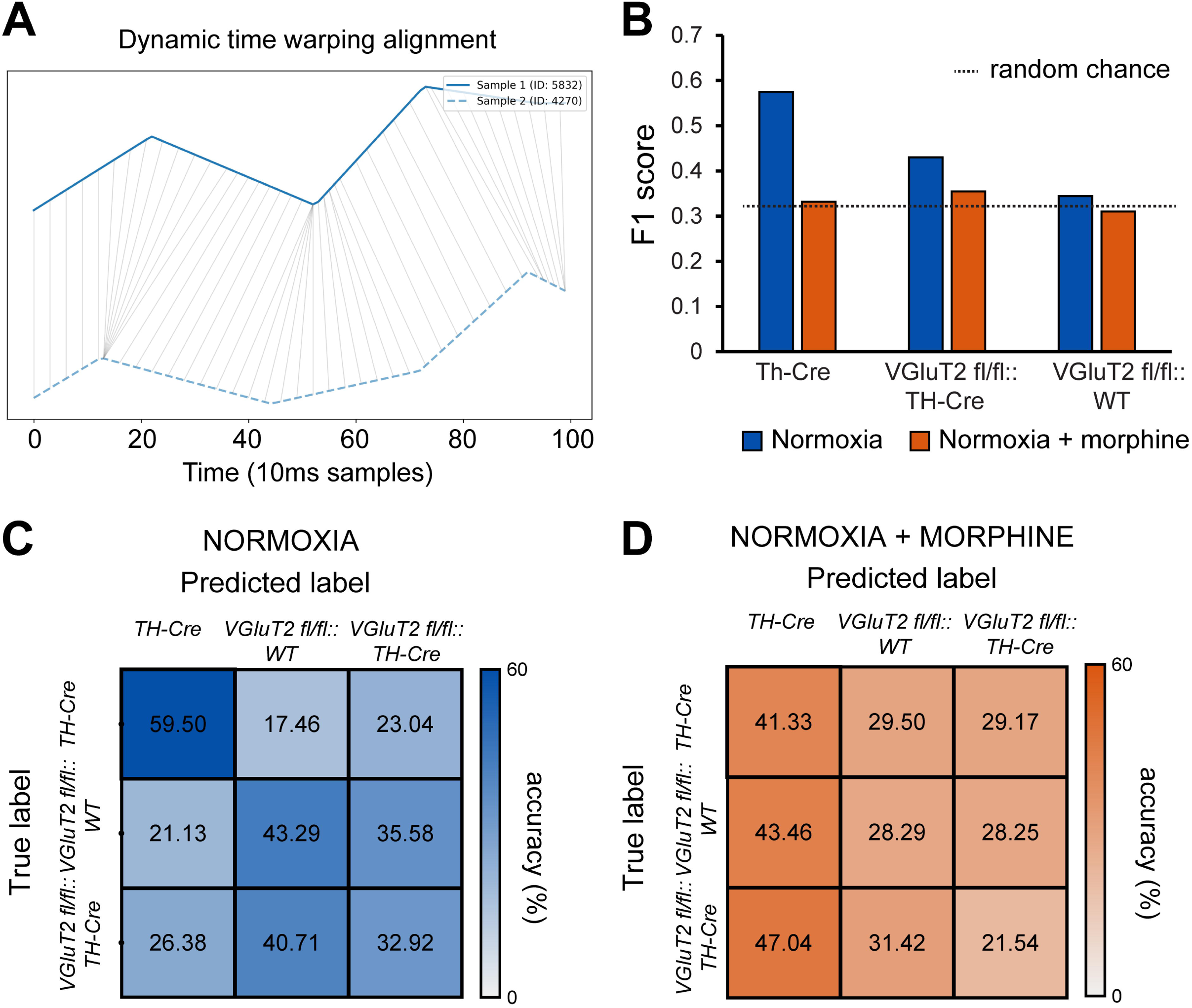
Machine learning applied to respiratory traces with DTW revealed genotype-dependent breathing pattern differences. (**A**) 1 sec-long instantaneous frequency traces (in 10 ms bins) were subjected to dynamic time warping (DTW) to assess the distance in proximal time-points. (**B**) The F1 score is used as the metric of model performance for breathing classification. Dashed line (33.3%) represents chance and poor classification performance. (**C**)-(**D**) Confusion matrices from DTW-kNN analyzed data demonstrating the classifier’s accuracy in predicting the subjects’ genotype based on respiration in normoxia (**C**) and following morphine injection in normoxia (**D**).

The classifier was trained and tested on instantaneous frequency time series data under two conditions: normoxia baseline and normoxia following morphine administration. During baseline normoxia, the F1 score (metric of model performance) indicates that the classifier successfully distinguished among the three genotypes at a rate significantly above random chance (>33.3%; **Fig 4B**). Classification performance, depicted by the confusion matrices in **Figure 4 C and D**, was strongest for the TH-Cre group (59.5% correct) and VGluT2fl/fl::TH-Cre group (43.3% correct), though the model showed substantial confusion between VGluT2fl/fl::WT and VGluT2fl/fl::TH-Cre groups (40.7% misclassification rate; **Fig 4C**). In contrast, during morphine-induced respiratory depression, classifier performance dropped to 30.4% mean accuracy - near or below chance level. The confusion matrix revealed extensive misclassifications across all groups, with a strong bias toward predicting the TH-Cre genotype (**Fig 4D**).

These results indicate that glutamate co-release from catecholaminergic neurons contributes to a characteristic temporal structure in the breathing pattern under baseline conditions. The collapse of this pattern discrimination during opioid exposure suggests that opioid-induced inhibition — likely both presynaptic suppression of glutamate release from LC and other catecholaminergic neuron terminals, and postsynaptic hyperpolarization of KF and other opioid-sensitive respiratory neurons (such as shown in **Fig 1**) — effectively eliminates the genotype-specific breathing dynamics observed at baseline.

## Discussion

Here, we demonstrate that glutamate co-release from catecholaminergic neurons contributes to the control of breathing and may be susceptible to opioid inhibition through both presynaptic and postsynaptic mechanisms. Using cell type-specific optogenetics, we show that in our representative circuit model, LC to KF glutamate co-release is potently inhibited by opioids, while excitatory NAergic signaling appears resistant (**Fig 1**). In line with these findings, freely behaving mice lacking VGluT2 in all TH+ neurons display altered respiratory patterning under baseline conditions and during hypercapnia compared to control TH-Cre hemizygous mice, with morphine administration diminishing these differences (**Fig 2-3**). Together, these data provide support for glutamate release from catecholaminergic neurons as a state-dependent respiratory modulator that is selectively vulnerable to opioid suppression.

Our results from brain slice recordings reveal that opioids inhibit glutamatergic LC-KF transmission through convergent pre- and postsynaptic mechanisms. Specifically, the opioid agonist ME reduced optically-evoked glutamatergic EPSC amplitudes and increased paired-pulse ratios, consistent with presynaptic opioid receptor-mediated inhibition of vesicle release from LC terminals (**Fig 1**). This finding aligns with the high density of MORs on LC neurons and their well-established role in suppressing LC activity (46, 47). Importantly, we also found that approximately half of the glutamate-modulated KF neurons exhibit GIRK currents in response to opioid exposure, indicating direct postsynaptic inhibition. This dual sensitivity creates a multiplicative inhibitory effect where opioids simultaneously reduce transmitter release and hyperpolarize some of the target neurons — a circuit architecture that may render the LC-KF glutamatergic pathway particularly vulnerable during OIRD.

In contrast, KF neurons excited by NAergic signaling show minimal opioid sensitivity, with only 12.5% of all NA-responsive KF neurons displaying GIRK currents, and no presynaptic impact on NA release from the LC. This partial opioid-inhibition complements our previous conclusion that co-released NA and glutamate are most likely packaged into independent vesicles at the LC axon terminal (18). Based on this, and the growing literature on neuromodulator and small molecule neurotransmitter co-release (48), the differential vulnerability suggests that glutamate and NA release from the LC may serve distinct functional roles in respiratory control, with the glutamatergic component providing state-dependent excitatory drive that is selectively suppressed by opioids. The apparent preservation of NA signaling during opioid exposure may partially contribute to the persistence of respiratory drive, albeit at depressed levels, during OIRD.

Further, the results of our plethysmography experiments in VGluT2^fl/fl^::TH-Cre mice support a specific modulatory role for glutamate co-release from catecholaminergic neurons in respiratory patterning. Under baseline conditions, these mice exhibit prolonged inspiratory duration, increased tidal volume, and reduced respiratory rate compared to TH-Cre controls, while maintaining similar minute ventilation (**Fig 2**). This reciprocal relationship between tidal volume and rate suggests that glutamate co-release likely promotes faster, shallower breathing patterns. Considering the general principles of glutamate co-release in other systems, for instance the low spiking-threshold and temporal precision of glutamate release (49), specific, time-locked respiratory modulation is likely.

Our findings should be considered within the broader context of recent work describing catecholaminergic contributions to respiratory control. Sun and Ray (11) demonstrated that chemogenetic inhibition of TH-Cre neurons disrupts hypercapnic and hypoxic chemoreflexes, establishing that catecholaminergic neurons are required for full chemosensory responses. More recently, Chang and colleagues showed that selective deletion of VGluT2 from DBH+ neurons — targeting NAergic but not dopaminergic populations — had no detectable effect on baseline breathing or chemoreflexes (29). Here, we deleted VGluT2 from all TH+ neurons (including both NAergic and dopaminergic populations) and observed alterations in respiratory patterning, linking breathing differences to both dopaminergic modulation and NAergic drive.

The contrast between these studies suggests several important conclusions. First, glutamatergic signaling from NAergic neurons alone may be insufficient to detectably influence baseline breathing (29), whereas glutamate co-release from the broader catecholaminergic network, including dopaminergic groups, does contribute to pattern generation. Second, catecholaminergic neurons can support chemoreflexes through neuromodulator signaling even when glutamate co-release is eliminated (29), consistent with Sun and Ray’s demonstration that inhibiting the entire TH+ population disrupts these reflexes (11). Third, our observation of altered pattern without abolished chemoreflexes indicates that glutamate release from catecholaminergic neurons plays a modulatory role in refining respiratory output rather than serving as an essential component of the basic respiratory control system or chemosensory pathways.

Specific genotype-dependent differences became more pronounced during hypercapnia, where VGluT2^fl/fl^::TH-Cre mice showed exaggerated increases in respiratory rate and corresponding decreases in tidal volume compared to controls (**Fig 3**). This suggests that glutamate release from catecholaminergic neurons may be particularly important for sculpting the hypercapnic response pattern, possibly by modulating the balance between relative changes in rate and tidal volume. Importantly, the hypercapnic ventilatory reflex remained intact in all groups, which is consistent with findings by Chang et al. showing that VGluT2-based glutamatergic signaling is not required for chemosensory reflexes (29). However, our results reveal that glutamate co-release from all catecholaminergic neurons does change the pattern of the hypercapnic response.

Following morphine administration, the differences between VGluT2-deficient and control mice were substantially reduced (**Fig 3**). The machine learning approach employing DTW reinforced this finding, successfully distinguishing between genotypes under baseline conditions but failing to do so after morphine (**Fig 4**). This convergence supports our circuit-level findings that opioids preferentially suppress glutamatergic transmission. When glutamate signaling is pharmacologically inhibited by morphine in control animals, their respiratory patterns become more similar to mice lacking VGluT2-dependent glutamate release from TH positive neurons. This pattern difference complements our demonstration of opioid-sensitive glutamatergic currents in the LC-KF circuit and suggests that the state-dependent modulatory role of catecholaminergic glutamate co-release is specifically vulnerable during opioid exposure.

The LC-KF circuit only represents one of several catecholaminergic respiratory pathways potentially contributing to the observed behavioral phenotypes. The KF is well-positioned to integrate LC glutamatergic input with other respiratory signals, and its known role in phase-switching and respiratory pattern modulation makes it a plausible substrate for the altered inspiratory-expiratory timing we observed (50–52). The substantial opioid inhibition we observed in the LC-KF glutamatergic pathway provides a potential mechanism for how the KF contributes to OIRD (53–55). Previous work established that the KF is critical for opioid-induced rate depression and pattern irregularity. Our findings suggest that loss of excitatory glutamatergic input to KF neurons, through both reduced LC terminal release and direct postsynaptic inhibition, may contribute to the KF’s inability to maintain normal phase-switching dynamics during opioid exposure. However, whether this specific LC-KF pathway mediates any of the observed behavioral effects of morphine remains to be tested through pathway-specific manipulations. Our brain slice experiments necessarily isolate the LC-KF circuit from the broader respiratory network, and the functional consequences of opioid inhibition at this synapse may differ in the intact system where multiple compensatory mechanisms will operate.

In addition to this hub, TH+ neurons also project to other respiratory nuclei, and dopaminergic populations may contribute through direct or indirect pathways independently. Our use of TH-Cre mice to delete VGluT2 encompasses multiple catecholaminergic populations that were not targeted in the DBH-Cre studies of Chang et al (29). The contrast between our findings and those of Chang et al. (29) highlights the need for intersectional genetic approaches to dissect the relative contributions of LC, A5, and other NAergic nuclei, as well as specific dopaminergic populations to respiratory pattern modulation. For instance, MORs are abundantly expressed in many dopaminergic nuclei (ventral tegmental area, substantia nigra, striatum), out of which the tubular striatum and the ventral tegmental area have been linked to some role in respiratory modulation (2, 38, 56–58).

In conclusion, we identified glutamate co-release from catecholaminergic neurons as a state-dependent modulator of respiratory patterning that is selectively vulnerable to opioid inhibition. By bridging the findings that TH+ neurons are required for full chemoreflexes (11), and that DBH+ glutamate signaling is dispensable for basic respiratory function (29), our results reveal a nuanced picture. It seems that glutamate co-release from the broader catecholaminergic network, and likely a dopaminergic source, refines respiratory patterns without being essential for basic respiratory patterning and the hypercapnic chemoreflex. The convergent pre- and postsynaptic opioid sensitivity of LC-KF glutamatergic transmission, combined with the diminished genotype differences after morphine administration, suggests that suppression of glutamate co-release from catecholaminergic neurons contributes to opioid-induced alterations in respiratory pattern. These findings advance our understanding of how catecholaminergic systems modulate breathing beyond classical monoaminergic mechanisms and highlight the circuit-level and synaptic complexity underlying OIRD.

## Acknowledgments

This research was supported by Thomas H. Maren Research Excellence Award to A. G. V., NIH grant K99/R00 HL159232 to A. G. V.

